# Combined computational and cellular screening identifies synergistic inhibition of SARS-CoV-2 by lenvatinib and remdesivir

**DOI:** 10.1101/2021.03.19.435806

**Authors:** Marie O. Pohl, Idoia Busnadiego, Francesco Marrafino, Lars Wiedmer, Annika Hunziker, Sonja Fernbach, Irina Glas, Elena V. Moroz-Omori, Benjamin G. Hale, Amedeo Caflisch, Silke Stertz

## Abstract

Rapid repurposing of existing drugs as new therapeutics for COVID-19 has been an important strategy in the management of disease severity during the ongoing SARS-CoV-2 pandemic. Here, we used high-throughput docking to screen 6000 compounds within the DrugBank library for their potential to bind and inhibit the SARS-CoV-2 3CL main protease, a chymotrypsin-like enzyme that is essential for viral replication. For 19 candidate hits, parallel *in vitro* fluorescence-based protease-inhibition assays and Vero-CCL81 cell-based SARS-CoV-2 replication-inhibition assays were performed. One hit, diclazuril (an investigational anti-protozoal compound), was validated as a SARS-CoV-2 3CL main protease inhibitor *in vitro* (IC50 value of 29 µM) and modestly inhibited SARS-CoV-2 replication in Vero-CCL81 cells. Another hit, lenvatinib (approved for use in humans as an anti-cancer treatment), could not be validated as a SARS-CoV-2 3CL main protease inhibitor *in vitro*, but serendipitously exhibited a striking functional synergy with the approved nucleoside analogue remdesivir to inhibit SARS-CoV-2 replication, albeit this was specific to Vero-CCL81 cells. Lenvatinib is a broadly-acting host receptor tyrosine kinase (RTK) inhibitor, but the synergistic effect with remdesivir was not observed with other approved RTK inhibitors (such as pazopanib or sunitinib), suggesting that the mechanism-of-action is independent of host RTKs. Furthermore, time-of-addition studies revealed that lenvatinib/remdesivir synergy probably targets SARS-CoV-2 replication subsequent to host-cell entry. Our work shows that combining computational and cellular screening is a means to identify existing drugs with repurposing potential as antiviral compounds. Future studies could be aimed at understanding and optimizing the lenvatinib/remdesivir synergistic mechanism as a therapeutic option.

## INTRODUCTION

Severe acute respiratory syndrome coronavirus 2 (SARS-CoV-2) entered the human population in late 2019 and rapidly spread around the globe causing a devastating pandemic with significant impacts on human health and the worldwide economy [1, 2]. Infection with SARS-CoV-2 results in a respiratory disease, COVID-19, that is mild in many individuals or even asymptomatic. However, in vulnerable patients, in particular individuals with compromised immune systems, such as the elderly, immunosuppressed or those with other chronic diseases, the virus can cause a severe life-threatening pneumonia [2-5]. Thus, large efforts have been undertaken to develop vaccines and antiviral drugs for the prevention and treatment of COVID-19. While several vaccines have been developed and licensed at unprecedented speeds, the development of antiviral drugs has not yet been as successful [6-9]. Remdesivir is the only antiviral drug that is currently approved and used to treat COVID-19 patients in many countries, but its efficacy has been questioned by recent results from a large-scale clinical trial [10, 11]. Thus, additional efforts are urgently required to fill this gap.

Early on in the pandemic, drug development approaches focused on the repurposing of drugs approved for other conditions as a means to accelerate the process of bringing drug candidates to market [12, 13]. One of the viral drug targets that has gained particular interest is the SARS-CoV-2 main protease, 3CL M^pro^, a chymotrypsin-like protease that cleaves the two polyproteins, pp1a and pp1ab, and is essential for viral replication [14, 15]. 3CL M^pro^ is considered a promising target for drug development as its structure has been solved and there is no homologous human protease [16, 17]. Moreover, protease inhibitors have been developed successfully to treat HIV-1 and hepatitis C virus infections [18, 19]. While several inhibitors of 3CL M^pro^ have already been identified, no drug targeting 3CL M^pro^ has been approved yet [16, 20-23]. Here, we computationally screened a library of approved drugs for novel 3CL M^pro^ inhibitors. The identified drug candidates were then tested for their antiviral potential in an *in vitro* 3CL M^pro^ protease-inhibition assay, and in a Vero cell-based virus replication assay using authentic SARS-CoV-2. Two potential inhibitors (diclazuril and ramelteon) identified *in silico* possessed modest antiviral activity. Furthermore, we found that lenvatinib, an inhibitor of receptor tyrosine kinases such as vascular endothelial growth factor receptors, displayed strong synergistic inhibition of SARS-CoV-2 with the approved drug remdesivir.

## RESULTS

To identify novel inhibitors of the SARS-CoV-2 3CL M^pro^, we performed an *in silico* screening campaign of the DrugBank library which contains approximately 6000 approved drugs [24]. The docking was carried out by the computer program SEED (details in Methods section) using two holo structures of 3CL M^pro^ (PDB codes 5r82 and 5re9). The binding pocket was considered rigid during docking. The calculation of the binding energy makes use of a force field-based methodology. Upon filtering and ranking, the top 19 compounds were purchased for experimental validation (Fig. 1A, B).

**Fig. 1.**
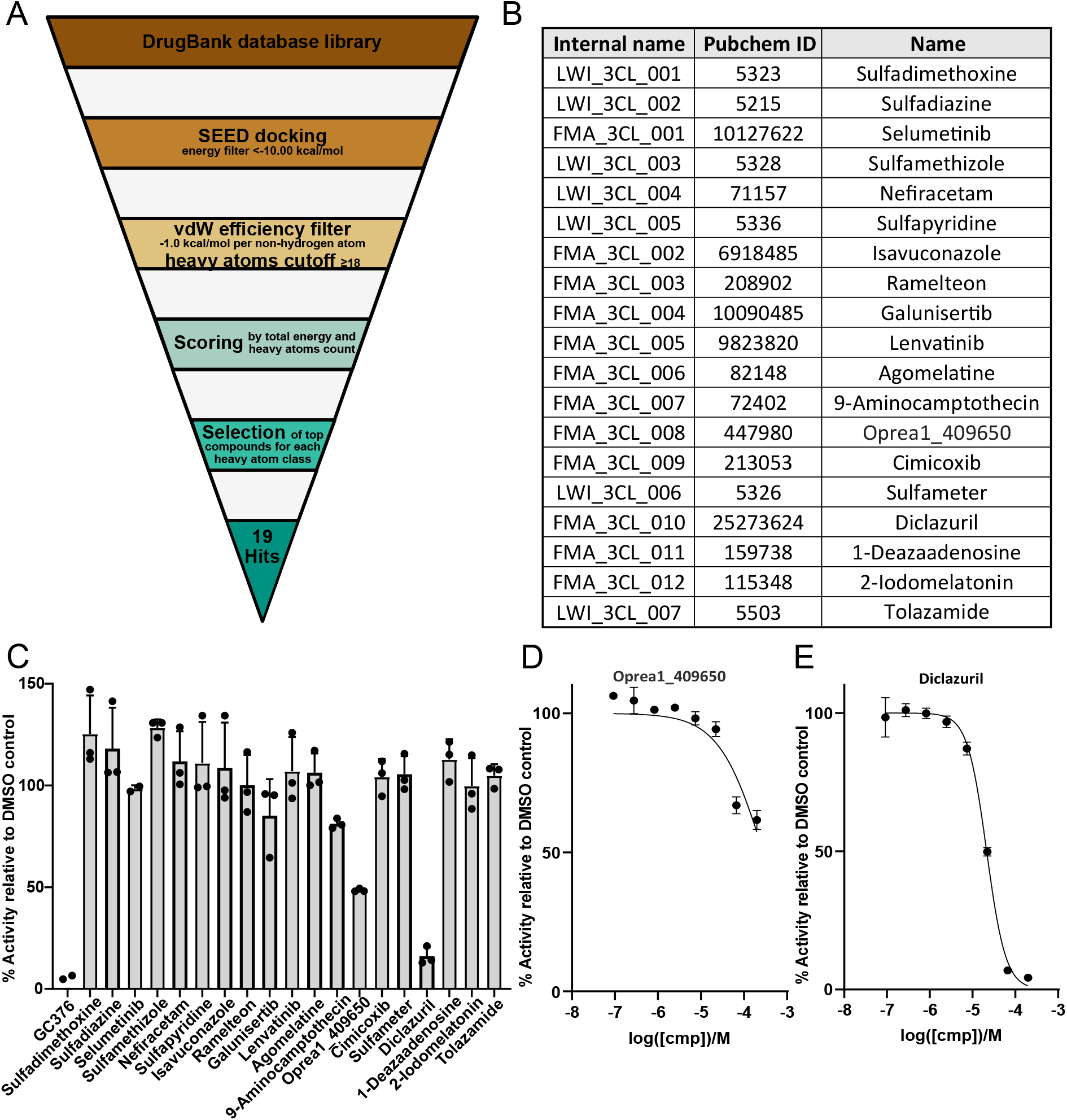
*In silico* screen reveals novel inhibitors of the SARS-CoV-2 3CL M^pro^ protease. **A)** Schematic overview of the *in silico* repurposing screen by high-throughput docking of known drugs. **B)** List of 19 compounds identified as potential inhibitors of 3CL M^pro^. **C)** The 19 compounds were tested in an *in vitro* fluorescence-based assay at a concentration of 100 µM for their potential to inhibit 3CL M^pro^. Mean values from three independent results are shown with error bars representing standard deviations. GC376 is the positive control. **D-E)** The two compounds that showed a reduction of enzymatic activity of more than 40% (relative to DMSO solution) in the *in vitro* protease assay, oprea1_409650 (D) and diclazuril (E), were tested at varying concentrations to obtain a dose-response curve. Average values from three independent experiments are plotted with error bars representing standard deviations.

An *in vitro* fluorescence-based protease-inhibition assay with recombinant 3CL M^pro^ protease was employed to measure the potential inhibition of enzymatic activity by each compound. The positive control, GC376, was used as reference compound [23]. An IC_50_ (inhibitor concentration that results in 50% reduction of activity with respect to buffer-only solution) value of 316 nM was measured for GC376, which is consistent with a reported value of 190 nM [23]. First, single-dose measurements were carried out at 100 µM for each compound (Fig. 1C). For only two compounds a substantial reduction (> 40% with respect to DMSO-only) of 3CL M^pro^ activity was measured. In follow-up dose-response experiments with these two compounds, diclazuril showed an IC_50_ of 29 µM, and oprea1_409650 showed an IC_50_ of 100 µM (Fig. 1D, E). The binding modes of diclazuril and oprea1_409650 as predicted by docking are shown in Fig. 2A-B. Both compounds are involved in hydrogen bonds with the backbone polar groups of Glu166 while they differ in the position and orientation of their phenyl rings (Fig. 2C).

**Fig. 2.**
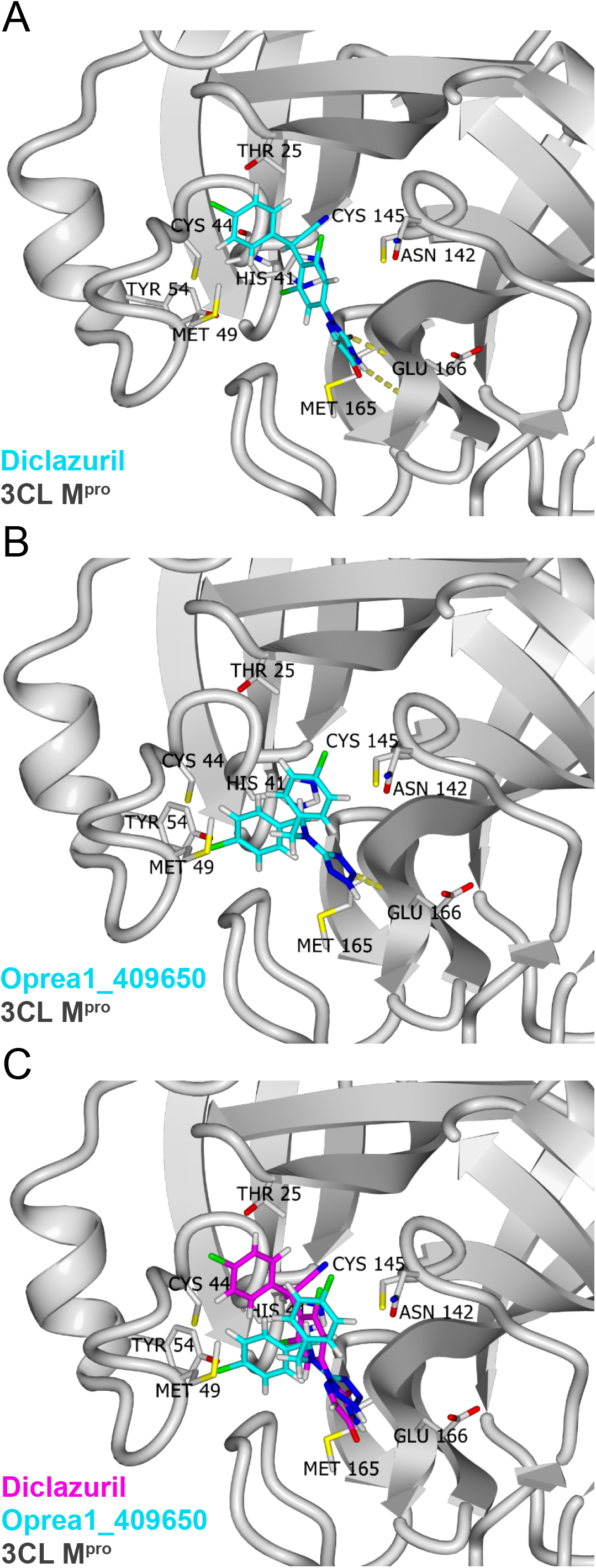
Predicted binding of diclazuril and oprea1_409650 to 3CL M^pro^. **A)** Predicted binding mode of diclazuril (carbon atoms in cyan) to 3CL M^pro^ (ribbon model in grey). The hydrogen bonds between diclazuril and the backbone polar groups of Glu166 are shown (yellow dotted lines). **B)** Predicted binding mode of oprea1_409650 (same colour scheme as in **A**). **C)** Comparison of the predicted binding mode of diclazuril (carbon atoms in magenta) and oprea1_409650 (carbon atoms in cyan). The Cα atoms of 3CL M^pro^ were used for the structural overlap.

In parallel, we tested the 19 predicted 3CL M^pro^ protease inhibitors for their potential to inhibit SARS-CoV-2 replication in Vero-CCL81 cells, an African Green Monkey kidney cell line shown to support productive SARS-CoV-2 replication [25]. Following pretreatment with the different inhibitors for 2 h, Vero-CCL81 cells were infected with SARS-CoV-2 isolate SARS-CoV-2/human/Switzerland/ZH-395-UZH-IMV5/2020 at a multiplicity of infection (MOI) of 0.01 PFU/cell and viral titers were determined at 24 h post-infection. Remdesivir (RDV), a known inhibitor of SARS-CoV-2 [26], was included as a positive control. In parallel, inhibitor-treated cells were left uninfected and cell viability was determined. With the exception of compounds 9-aminocampthecin and 1-deazaadenosine, cell viability was only minimally affected by treatment with the inhibitors (Fig. 3A). In line with previous reports, SARS-CoV-2 replication was inhibited by remdesivir in a dose-dependent manner, thus validating our screening assay (Fig. 3B). Interestingly, the compound selumetinib, a kinase inhibitor with specificity for mitogen-activated protein kinase kinase (MAPK kinase) subtypes 1 and 2, displayed a positive Z-score and lead on average to a 3.6-fold increase in viral titer despite reducing cell viability by 20% (Fig. 3A-B). However, as we aimed to identify viral inhibitors we focused on the compounds with negative Z-scores. We observed significant inhibition of virus replication with three of the predicted protease inhibitors, lenvatinib, ramelteon and diclazuril (Fig. 3B). Of note, diclazuril, which was the most potent protease inhibitor in the *in vitro* protease assay, was also the strongest hit in the screen for inhibition of SARS-CoV-2 replication. Oprea1_409650, which also displayed inhibition in the protease assay, did not reduce viral replication significantly. While the inhibitory effects of the three hit compounds were robust and reproducible across the replicates of the SARS-CoV-2 screening assay, the degree of inhibition was only around 50-60% and thus modest (Fig. 3C).

**Fig. 3.**
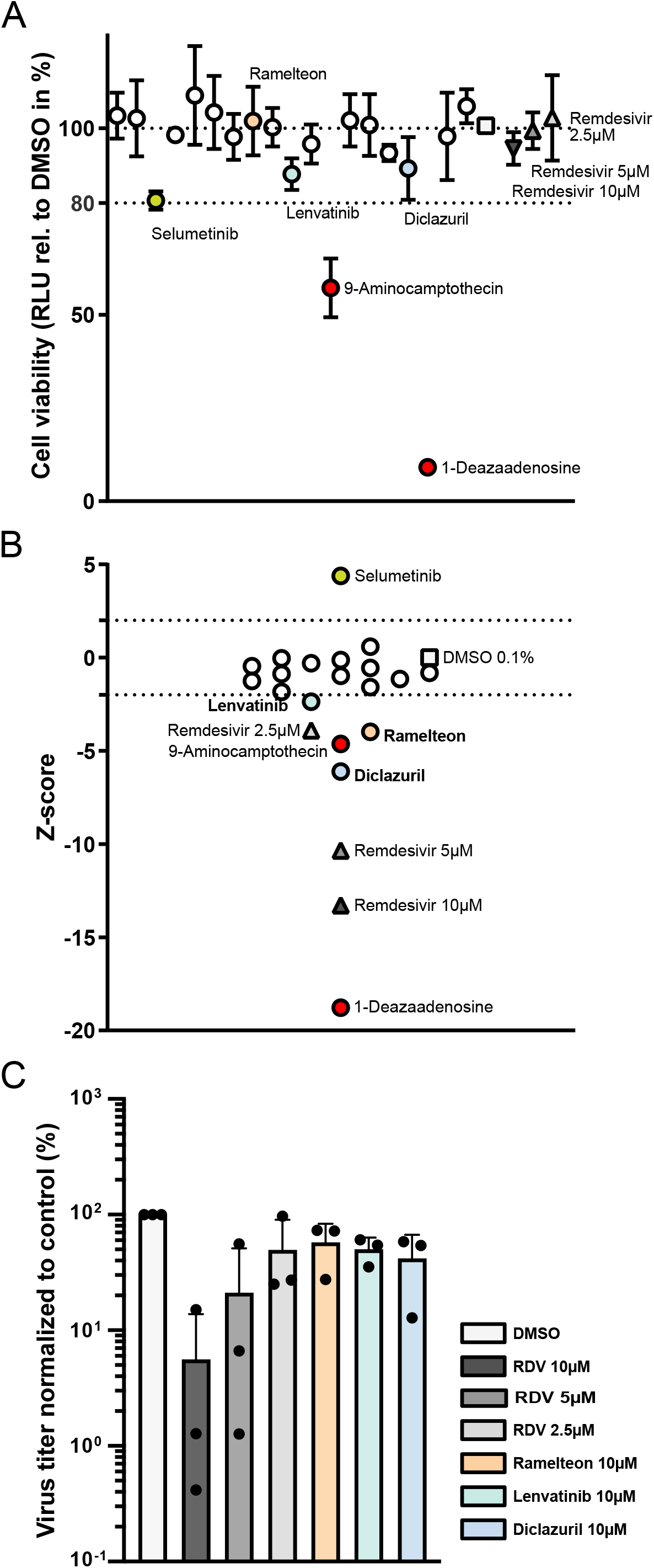
Vero-CCL81-based screen reveals diclazuril, lenvatinib and ramelteon as inhibitors of SARS-CoV-2. **A) Effect of inhibitor-treatment on cell viability**. Vero-CCL81 cells were pre-treated for 2 h with 10 µM of each putative 3CL M^pro^ inhibitor or with DMSO 0.1% or RDV (10, 5, and 2.5 µM) as negative and positive controls, respectively. Then, cells were mock-infected for 1 h followed by incubation in the presence of the respective inhibitors or controls. 24 h after mock-infection cell viability was assessed. Shown are means and standard deviation of cell viability normalized to the DMSO control (depicted as a square) of three independent experiments each performed in duplicates. RDV-treated cells are depicted as triangles. Compounds reducing cell viability more than 20% compared to the DMSO control (marked in red) were excluded from the selection. **B) Effect of inhibitor-treatment on SARS-CoV-2 growth**. Experimental set-up as in A) but cells were infected with SARS-CoV2 (MOI=0.01 PFU/cell) for 1 h followed by incubation in the presence of the respective inhibitors or controls. After 24 h of incubation, cell supernatants were harvested and viral growth was assessed by plaque assay. Viral titers of three independent experiments, each performed in duplicates, were normalized to the DMSO control, log2-transformed and converted into Z-scores. Z-score of DMSO and RDV-treated cells are depicted as a square and triangles, respectively. Excluded inhibitors are marked in red. A Z-score of smaller than -2 was used for the selection of inhibitors for follow-up. **C) Normalized virus titers for selected inhibitors**. Experimental set-up as in B). Shown are virus titers normalized to DMSO control of the selected inhibitors from B). Depicted are means and standard deviations of three independent experiments, each performed in duplicates.

To evaluate potential synergistic effects with an established inhibitor of SARS-CoV-2, we tested the predicted inhibitors of the protease in combination with low concentrations of remdesivir, which acts on a different target (the viral RNA-dependent RNA polymerase). While ramelteon and diclazuril did not increase virus inhibition by remdesivir, the combination of remdesivir and lenvatinib showed a very strong synergistic effect at 24 and 48 h post infection: the low concentrations of remdesivir used reduced viral titers by a maximum of 3-fold, but in combination with lenvatinib, SARS-CoV-2 titers were reduced approximately 1000-fold (Fig. 4A-B), which was not due to cytotoxic effects (Fig. 4C-D). When the three hit compounds were tested in human Calu-3 cells we did not observe inhibition of virus growth, either by the inhibitors alone or in combination with low doses of remdesivir (Fig. 4E). When we tested the combination of a low dose of remdesivir together with lenvatinib for the potential to inhibit SARS-CoV-2 in primary human nasal epithelial primary cultures (NEpC), we observed strong inhibition of virus replication in two out of three independent replicates. However, as the third experiment did not confirm this effect, we cannot conclude that there is a robust synergistic effect of these compounds in human NEpCs (Fig. 4F). Thus, the striking synergistic effect of lenvatinib in combination with remdesivir appears to be specific to Vero-CCL81 cells.

**Fig. 4.**
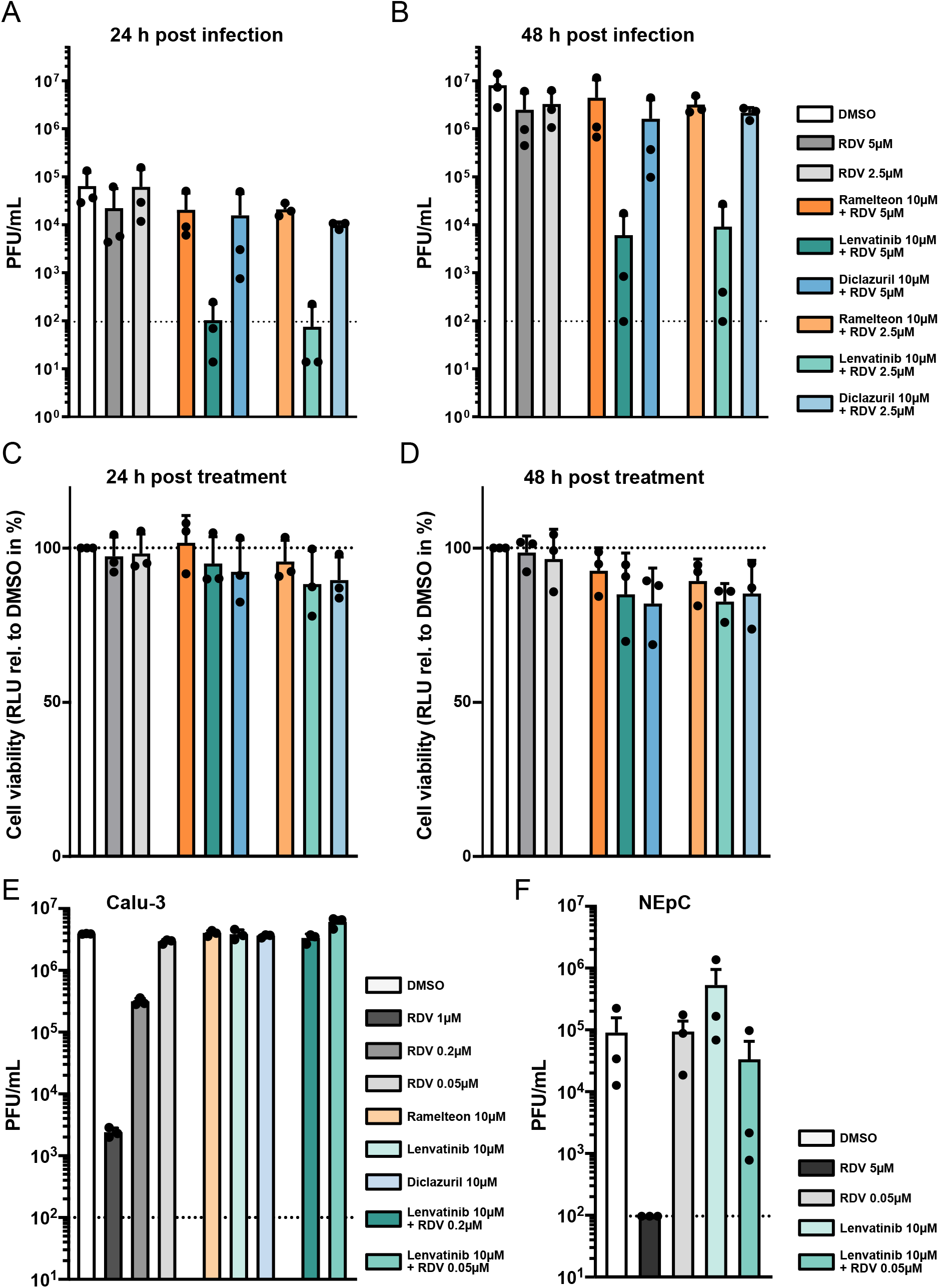
Lenvatinib acts synergistically with remdesivir to inhibit SARS-CoV-2. **A-B) Effect of combining selected inhibitors with RDV on SARS-CoV-2 growth**. Vero-CCL81 cells were pre-treated for 2 h with 10 µM of ramelteon, lenvatinib, or diclazuril combined with either 5 or 2.5 µM RDV. DMSO 0.1% or RDV (5 and 2.5 µM) were used as negative and positive controls, respectively. Following infection with SARS-CoV-2 (MOI=0.01 PFU/cell) for 1 h, cells were incubated in the presence of the respective inhibitors or controls. After 24 h (A) and 48 h (B) of incubation, cell supernatants were harvested and viral growth was assessed by plaque assay. Shown are mean virus titers and standard deviations of three independent experiments, each performed in duplicates. Dashed line indicates limit of detection. **C-D) Effect of combining selected inhibitors with RDV on Vero-CCL81 cell viability**. Experimental set-up as in A-B, but cells were mock-infected and cell viability was assessed at 24 h (C) or 48 h (D) post mock infection. Shown are means and standard deviations of cell viability normalized to the DMSO control of three independent experiments each performed in duplicate. A-D) The colour code for the different bars in A-D is shown next to panel B. **E) Effect of combination of selected inhibitors with RDV on SARS-CoV-2 viral growth in Calu-3 cells**. Calu-3 cells were pre-treated for 2 h with 10 µM of ramelteon, lenvatinib or diclazuril; or 10 µM lenvatinib combined with either 0.2 or 0.05 µM RDV. DMSO 0.1% or RDV (1, 0.2 and 0.05 µM) were used as negative and positive controls, respectively. Cells were then infected for 1 h with SARS-CoV-2 (MOI=0.1 PFU/cell) followed by incubation in the presence of the respective inhibitors or controls. After 24 h of incubation, cell supernatants were harvested, and viral growth was assessed by plaque assay. Shown are mean virus titers and standard deviations of three independent experiments, each performed in duplicates. Dashed line indicates limit of detection. **F) Effect of combination of lenvatinib with RDV on SARS-CoV-2 viral growth in primary nasal epithelial cultures (NEpCs)**. NEpCs were pre-treated for 2 h with 10 µM of lenvatinib or 10 µM lenvatinib combined with 0.05 µM RDV. DMSO 0.1% or RDV (5 and 0.05 µM) were used as controls. Cells were then infected for 1 h with SARS-CoV-2 (6×10^4^ PFU/well) followed by incubation in the presence of the respective inhibitors or controls. After 48 h of incubation, cell supernatants were harvested, and viral growth was assessed by plaque assay. Shown are mean virus titers and standard deviations of three independent experiments. Dashed line indicates limit of detection.

As lenvatinib did not inhibit SARS-CoV-2 3CL M^pro^ in the *in vitro* protease-inhibitor assay at the concentration used in cell culture-based assays (Fig. 1C), we speculated that its function as an inhibitor of receptor tyrosine kinases (RTKs) might contribute to the observed synergy with remdesivir. We therefore selected pazopanib and sunitinib as additional RTK inhibitors to test for synergy with remdesivir, as they exhibit a similar inhibition profile to lenvatinib, with all three shown to inhibit vascular endothelial growth factor receptors (VEGFR1, VEGFR2 and VEGFR3), platelet-derived growth factor receptor (PDGFR-α/-β), and c-Kit, with comparable specificity and potency [27, 28]. When taking cell viability into consideration, only lenvatinib displayed a clear synergistic effect with remdesivir (Fig. 5A-B) suggesting that the observed synergistic inhibitory effect is not linked to inhibition of VEGFR-, PDGFR- or c-Kit induced signaling. Next, we performed a time-of-addition experiment, in which we treated cells with remdesivir, lenvatinib, or the combination of both inhibitors, at 1 h prior to infection, 1 h post infection, or 4 h post infection. At 24 h post infection, supernatants were harvested and infectious virus titers determined by plaque assay. As expected from an RNA polymerase inhibitor, remdesivir retained its antiviral potential if given post viral entry: treatment with remdesivir 1 h post infection resulted in a similar inhibition of virus replication as treatment of cells prior to infection (Fig. 5C). In addition, treatment with remdesivir at 4 h post infection still led to a substantial decrease in viral replication. The synergistic effect of lenvatinib and remdesivir was highly similar for pretreatment and 1 h post-treatment conditions, both resulting in potent inhibition of virus replication. Furthermore, the 4 h post infection treatment condition resulted in a less potent, but still substantial, decrease in viral titers by more than 10-fold (Fig. 5C). When we assessed expression of the viral nucleocapsid protein, N, at 7 h post infection we again saw that the combination of lenvatinib and remdesivir potently inhibited SARS-CoV-2 (Fig. 5D). In line with the results from the time-of-addition experiment, we observed a strong reduction in signal intensity for the N staining rather than a reduction in the number of infected cells (Fig. 5D). Thus, our data suggest that the synergistic effect of lenvatinib and remdesivir could be due to more potent inhibition of the viral transcription and/or replication processes rather than any effects on earlier stages of the viral replication cycle.

**Fig. 5.**
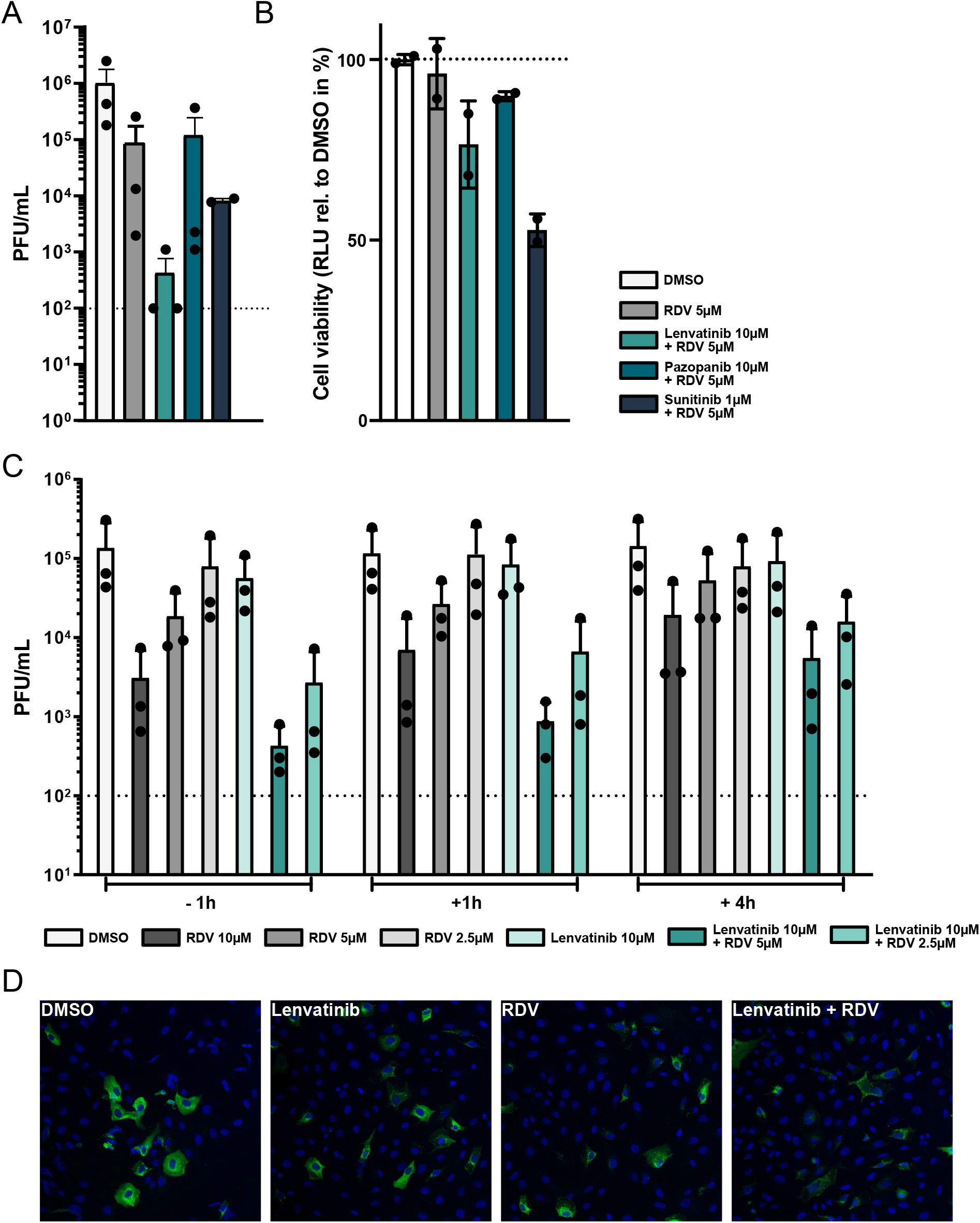
The combination of lenvatinib and remdesivir inhibits SARS-CoV-2 at a post-entry step. **A) Effect of VEGFR inhibitors and RDV on SARS-CoV-2 growth**. Vero-CCL81 cells were pre-treated for 2 h with lenvatinib (10 µM), pazopanib (10 µM), or sunitinib (1 µM) in combination with RDV (5 µM). DMSO 0.1% or RDV (5 µM) were used as negative and positive controls, respectively. Following infection with SARS-CoV2 (MOI=0.01 PFU/cell) for 1 h, cells were incubated in the presence of the respective inhibitors or controls. After 48 h of incubation, cell supernatants were harvested and viral growth was assessed by plaque assay. Shown are mean virus titers and standard deviations of three independent experiments, each performed in duplicates. Dashed line indicates limit of detection. **B) Effect of VEGFR inhibitors and RDV on Vero-CCL81 cell viability**. Experimental set-up as in A, but cells were mock-infected. 48 h after mock-infection, cell viability was assessed. Shown are means and standard deviations of cell viability normalized to the DMSO control of two independent experiments each performed in duplicate. **C) Effect of time-of-addition on the synergistic reduction of SARS-CoV-2 growth by lenvatinib and RDV**. Vero-CCL81 cells were treated either 1 h before (−1h), 1 h after (+1h) or 4 h after (+4h) infection with SARS-CoV-2 (MOI=0.01 PFU/cell) with 10 µM of lenvatinib combined with either 5 or 2.5 µM RDV. DMSO 0.1% or RDV (10, 5 and 2.5 µM) were used as negative and positive control, respectively. 24 h post infection, cell supernatants were harvested and viral growth was assessed by plaque assay. Shown are mean virus titers and standard deviations of three independent experiments, each performed in duplicates. Dashed line indicates limit of detection. **D) Inhibition of viral N protein expression by lenvatinib and remdesivir**. Vero-CCL81 cells were pre-treated for 2 h with lenvatinib (10 µM), remdesivir (2.5 µM), or the combination of both (10 µM lenvatinib + 2.5 µM remdesivir). DMSO was used as negative control. Following infection with SARS-CoV2 (MOI=5 PFU/cell) for 7 h in the presence of the inhibitors, cells were fixed and stained for SARS-CoV-2 N (green) and nuclei (blue).

## DISCUSSION

Our approach of combining *in silico* and *in vitro* screenings of an approved drug library for inhibitors of the SARS-CoV-2 3CL M^pro^ protease identified diclazuril as a low micromolar inhibitor of the 3CL M^pro^ protease, and three drugs (diclazuril, ramelteon, and lenvatinib) that reduced viral replication in Vero-CCL81 cells. Moreover, lenvatinib showed an unexpectedly strong synergy with remdesivir. This shows that the approach used here can identify compounds with antiviral potential.

There are several caveats to the approach adopted in this study. Of note, we only tested the compounds at a single 10 µM concentration, and thus an immediate option to see stronger inhibition of virus replication would be to test higher concentrations of the inhibitors. However, this might result in the identification of additional compounds with antiviral potential but only in a non-physiologically achievable range of concentrations. Thus, extensive compound modification and optimization for increased antiviral potential would be required in subsequent studies and would slow immediate roll-out of a repurposed drug. Given that our goal was to rapidly identify approved drugs as treatment options for COVID-19 and circumvent the lengthy process of progressing inhibitors to approval, we set the threshold at 10 µM and did not consider higher drug concentrations. Options to improve the yield of our screening approach could be to use an alternative structure of 3CL M^pro^ that has subsequently been solved by room temperature X-ray crystallography [29]. It reveals the ligand-free structure of the active site and the conformation at near-physiological temperature, and was suggested to be a better target for *in silico* screening for inhibitors of 3CL M^pro^ [29]. Moreover, additional libraries of approved drugs could be included, for example IUPHAR/BPS (https://www.guidetopharmacology.org/) and ChEMBL (https://www.ebi.ac.uk/chembl/), although they overlap substantially with DrugBank.

A second caveat is the apparent cell-type specificity of synergy observed between lenvatinib and remdesivir: this effect was clear in Vero-CCL81 cells, but was not evident in human Calu-3 cells, or robust in human NEpCs. One possible explanation could be that lenvatinib inhibits an African green monkey cell-specific target that is not inhibited in human cells. This could either be due to differential expression levels of the respective target, or could be related to differences in target sequence and structure between the human and the African green monkey version of the target. As the conversion of remdesivir to its active metabolite (remdesivir triphosphate) is less efficient in Vero-CCL81 cells as compared to human cells [26], the synergistic effect of lenvatinib with remdesivir could be linked to the metabolic conversion of remdesivir. If lenvatinib treatment led to a higher conversion rate, and thus to higher levels of the active metabolite of remdesivir, it could be hypothesized that in human cells, where the conversion of remdesivir to remdesivir triphosphate is already efficient, an increase in conversion efficiency would not result in a boost of the antiviral potential of remdesivir. However, it is thus far unclear how lenvatinib would affect the metabolic conversion of remdesivir. Given that lenvatinib did not significantly inhibit the SARS-CoV-2 3CL M^pro^ protease in the *in vitro* protease assay, the known targets of lenvatinib (i.e. receptor tyrosine kinases (RTKs), such as VEGFR, PDGFR or c-Kit), could be the relevant drug targets for inhibition of virus replication and synergy with remdesivir. We did not observe synergy for two other RTK inhibitors with comparable RTK specificity and potency, and thus the observed synergy is probably not mediated by inhibition of VEGFRs, PDGFRs or c-Kit, which are targeted by all three inhibitors. Thus, future work should aim to reveal the mechanism behind the synergistic action of remdesivir and lenvatinib. Such studies on the Vero cell-specific synergy could reveal broader insights into the metabolism and antiviral mechanism of nucleoside analogs and thus aid the development of antiviral drugs.

## MATERIALS AND METHODS

### *In silico* screen for 3CL M^pro^ inhibitors

The evaluation of the binding energy by the program SEED makes use of a force field-based energy function. Protein and ligand electrostatic desolvation penalties are approximated by a continuum dielectric treatment and the generalized Born model [30-32]. The partial charges and vdW parameters for the atoms in the protein and in the ligands were taken from the CHARMM36 all-atom force field and the CHARMM general force field (CGenFF), respectively [33-35]. Importantly, CHARMM36 and CGenFF employ a consistent paradigm for the determination of partial charges and vdW parameters. The dielectric discontinuity surface was approximated by the molecular surface. The dielectric constant was assigned a value of 2.0 and 78.5 for the volume occupied by the solute and solvent, respectively. Default SEED parameters were used. The nearly 6000 approved and experimental drugs in DrugBank with more than 17 non-hydrogen atoms were docked by SEED. This filter on molecular sizes was employed to avoid the selection of very small molecules which cannot provide sufficient affinity. SEED offers three methods for the actual placement of the fragments: (1) polar docking in which fragments are positioned and oriented such that at least one favorable hydrogen bond with a residue in the binding site is formed, (2) apolar docking in which fragments are placed in the hydrophobic regions of the binding site, and (3) a combination of (1) and (2). The latter placement procedure was used. The docking was carried out on two holo structures of 3CL M^pro^ (PDB codes 5r82 and 5re9) with high resolution. The residue Glu166 was selected for the definition of the binding pocket. For PDB 5r82 no water molecules were retained, while for PDB 5re9 one buried water molecule was retained. Two energy filters were used: total binding energy < -10 kcal/mol and van der Waals efficiency < -1.0 kcal/mol per non-hydrogen atom. Hydrogen bonding penalty was calculated for each compound, according to the method described by Zhao and Huang [36]. If newly formed hydrogen bonds between the ligand and the protein are not in good geometry or desolvated donors/acceptors are not involved into new hydrogen bonds, an enthalpic loss arises due to uncompensated breakage of hydrogen bonds with water molecules. The hydrogen bonding penalty index is calculated by checking both the solvation and hydrogen bonding state of donors/acceptors at the protein-ligand interface. A value of 1 in penalty corresponds to about 1.5 kcal/mol in binding free energy. This penalty can then be integrated into total binding energy values according to the formula SEED total energy + hydrogen bonding penalty*1.5kcal/mol. Compounds were then ranked by recalculated total energy and heavy atom count. Up to five top compounds for each heavy atom count tier were selected and experimental phase drugs were removed, focusing on approved drugs. The resulting compounds were visually inspected, favoring compounds that established H-bonds, and 19 compounds in total were selected.

### *In vitro* protease inhibition assay

The *in vitro* inhibition assay was based on fluorescence, and was carried out according to the specifications of the vendor (Reaction Biology, M^pro^). The assay measures the inhibitory activity of compounds that compete with a known peptide substrate for purified 3CL M^pro^. The peptide contains a quencher and a donor group, thus fluorescence is quenched in an intact peptide. Upon cleavage of the peptide by 3CL M^pro^ the donor fragment generates fluorescence which is quantified by a luminometer. The fluorogenic substrate used is [NH2-C(EDANS)VNSTQSGLRK(DABCYL)M-COOH]. The positive control is GC376 which was employed as a reference compound. Single-dose screenings were performed at 100 µM.

### Compound screening for inhibitors of SARS-CoV-2

To test the selected compounds from the docking analysis for antiviral activity against SARS-CoV-2, 1.2×10^4^ Vero-CCL81 cells (ATCC) were seeded in DMEM supplemented with 10% fetal calf serum (FCS), 100 U/ml of penicillin, and 100□μg/ml of streptomycin in 96-well plates and incubated overnight at 37°C and 5% CO_2_. For the infection and cell viability experiments, compound dilutions of 10 µM and a final DMSO concentration of 0.1% were used. As negative control, 0.1% DMSO was used while 10, 5 and 2.5 µM remdesivir (at a final DMSO concentration of 0.1%) served as positive controls. The day after seeding, cells were washed with PBS followed by incubation with the individual compounds at a concentration of 10 µM diluted in Opti-MEM (Gibco) for 2 h at 37°C and 5% CO_2_. Next, medium was removed and cells were infected with SARS-CoV-2 (SARS-CoV-2/human/Switzerland/ZH-395-UZH-IMV5/2020) at a multiplicity of infection (MOI) of 0.01 PFU/cell diluted in PBS supplemented with 0.3% BSA, 1□mM Ca^2+^/Mg^2+^, 100 U/ml penicillin, and 100□μg/ml streptomycin [37]. Cells were then incubated for 1 h at 37°C and 5% CO_2_. Following infection, the inoculum was removed and DMEM supplemented with 10 µM of the respective compound (or the positive or negative control), 100 U/ml penicillin, 100□μg/ml streptomycin, 0.3% BSA, 20□mM HEPES, 0.1% FCS, and 0.5□μg/ml TPCK-treated trypsin was added to the cells. After incubation for 24 h at 37°C and 5% CO_2_, cell supernatants were harvested and virus titer was determined by plaque assay.

In parallel to the infection experiments, cell viability following incubation of Vero-CCL81 cells with the respective compounds was assessed using the same experimental set-up as for the infection experiments, but instead of infection with SARS-CoV-2, cells were mock-infected with PBS supplemented with 0.3% BSA, 1□mM Ca^2+^/Mg^2+^, 100 U/ml penicillin, and 100□μg/ml streptomycin for 1 h at 37°C and 5% CO_2_. Cell viability was determined 24 h after the addition of compound-containing DMEM using the CellTiter-Glo kit (Promega) and a luminescence plate reader (Perkin Elmer). The infection and cell viability experiments for the compound screen were performed in duplicates in three independent experiments.

For the analysis of the compound screen, all titers were normalized to the negative control (0.1% DMSO) and log2-transformed. Next, the Z-score was calculated by dividing the difference of the normalized and transformed titer of a given sample and the average normalized and transformed titer of the negative control by the standard deviation of the normalized and transformed negative control [38]. The average Z-score of the three independent replicates for a given sample was used for hit selection. Compounds associated with a Z-score of smaller than -2 were considered as hits for further analysis. Compounds associated with an average reduction in cell viability of more than 20% relative to the negative control were not considered as hits.

### Testing for synergistic inhibition of SARS-CoV-2 in Vero-CCL81 or Calu-3 cells

Vero-CCL81 or Calu-3 cells were seeded in 96 well plates and incubated overnight. Cells were washed once with PBS and incubated for 2 h at 37°C and 5% CO_2_ with Opti-MEM (Gibco) containing 5 or 2.5 μM remdesivir alone or in combination with 10 μM ramelteon (PubChem ID 208902), lenvatinib (PubChem ID 9823820), diclazuril (PubChem ID 25273624), pazopanib (LC laboratories, P-6706) or sunitinib (Selleckchem, S1042) at a final concentration of 0.1 % DMSO. DMSO 0.1 % was used as negative control. Cells were infected with SARS-CoV-2 (SARS-CoV-2/human/Switzerland/ZH-395-UZH-IMV5/2020) at a MOI of 0.01 PFU/cell in PBS supplemented with 0.3% BSA, 1 mM Ca^2+^/Mg^2+^, 100 U/ml penicillin, 100 μg/ml streptomycin. After 1 h incubation at 37°C and 5% CO_2_, cells were washed once in PBS and the indicated compounds were added. Samples were taken at 24 and 48 h post infection and supernatants were stored at -80°C prior to titer determination by plaque assay. In parallel to the infection experiments, cell viability was assessed as indicated in the compound screen section above. All experiments were performed in duplicates in three independent experiments.

### Testing for synergistic inhibition of SARS-CoV-2 in primary human nasal epithelial cultures

Primary human nasal epithelial cells (NEpCs) from a 41 year-old female donor were purchased from Epithelix (#EP51AB). Cells were grown in airway epithelium basal growth medium (Promocell, #C-21260) supplemented with an airway growth medium supplement pack (#C-39160; Promocell) and 10 µM Y-27632 (#1254; Tocris). Differentiation of NEpCs, and subsequent validation of airway cultures by measuring the transepithelial electrical resistance (TEER) was performed exactly as recently described [37]. NEpCs were washed once with PBS from the basal side and incubated for 2 h at 37°C and 5% CO_2_ with Gray’s medium containing the indicated inhibitors. Cells were then washed twice on the apical side before being infected with 6×10^4^ PFU of SARS-CoV-2 (SARS-CoV-2/human/Switzerland/ZH-395-UZH-IMV5/2020) in PBS supplemented with 0.3% BSA, 1 mM Ca^2+^/Mg^2+^, 100 U/ml penicillin, 100 μ g/ml streptomycin. After 1 h at 37°C and 5% CO_2_, the inoculum was taken off. Samples were taken at 48 h post infection and stored at -80°C until titer determination by plaque assay. Data represent mean values from three independent experiments with error bars depicting standard deviation.

### Time-of-addition experiments

Vero-CCL81 cells were seeded in 96 well plates at a density of 1.2 × 10^4^ cells/well. After incubation overnight, cells were washed once with PBS and incubated for 1 h at 37°C and 5% CO_2_ with Opti-MEM (Gibco) containing the indicated inhibitors. DMSO 0.1 % was used as negative control. Cells were infected with SARS-CoV-2 (SARS-CoV-2/human/Switzerland/ZH-395-UZH-IMV5/2020) at a MOI of 0.01 PFU/cell in PBS supplemented with 0.3% BSA, 1 mM Ca^2+^/Mg^2+^, 100 U/ml penicillin, 100 μg/ml streptomycin. Following infection, the inoculum was removed and DMEM supplemented with 100 U/ml penicillin, 100□μg/ml streptomycin, 0.3% BSA, 20□mM HEPES, 0.1% FCS and 0.5□μg/ml TPCK-treated trypsin was added to the cells. To test for time-of-addition effect, the indicated compounds were added either 1 or 4 h post infection instead of 1 h before infection in parallel samples. After incubation for 24 h at 37°C and 5% CO_2_, cell supernatants were harvested, and virus titer was determined by plaque assay. In parallel to the infection experiments, cell viability was assessed as indicated above. All experiments were performed in duplicates in three independent experiments.

### Immunofluorescence

Vero-CCL81 cells were seeded onto glass coverslips in 24 well plates at a density of 1 × 10^5^ cells/well and incubated overnight. Cells were washed once with PBS and incubated for 2 h at 37°C and 5% CO_2_ with Opti-MEM containing 2.5 μM remdesivir, 10 μM lenvatinib or a combination of 10 μM lenvatinib with 2.5 μM remdesivir. DMSO was used as negative control. Cells were infected with SARS-CoV-2 (SARS-CoV-2/human/Switzerland/ZH-395-UZH-IMV5/2020) at a MOI of 5 PFU/cell. After 1 h at 37°C and 5% CO_2_, cells were washed once in PBS and then incubated with inhibitor-containing medium for another 6 h. At 7 h post infection cells were fixed with 4% paraformaldehyde for 15 min, washed with PBS and permeabilized with PBS supplemented with 50 mM ammonium chloride (#254134; Sigma-Aldrich), 0.1% saponin (#47036, Sigma-Aldrich) and 2% BSA (#A7906; Sigma-Aldrich). Cells were stained with mouse anti-SARS-CoV-2 N (#MA5-29981; Thermo Fisher Scientific) and anti-mouse IgG Alexa488 (#A-11029; Thermo Fisher Scientific) as secondary antibody. Nuclei were stained with DAPI (#10236276001; Sigma-Aldrich).

